# Noise reduction strategies in metagenomic chromosome confirmation capture to link antibiotic resistance genes to microbial hosts

**DOI:** 10.1101/2022.11.05.514866

**Authors:** Gregory E. McCallum, Amanda E. Rossiter, Mohammed Nabil Quraishi, Tariq H. Iqbal, Sarah A. Kuehne, Willem van Schaik

## Abstract

The gut microbiota is a reservoir for antimicrobial resistance genes (ARGs). With current sequencing methods, it is difficult to assign ARGs to their microbial hosts, particularly if these ARGs are located on plasmids. Metagenomic chromosome conformation capture approaches (meta3C and Hi-C), have recently been developed to link bacterial genes to phylogenetic markers, thus potentially allowing the assignment of ARGs to their hosts on a microbiome-wide scale.

Here, we generated a meta3C dataset of a human stool sample and used previously published meta3C and Hi-C datasets to investigate bacterial hosts of ARGs in the human gut microbiome. Sequence reads mapping to repetitive elements were found to cause problematic noise in, and may importantly skew interpretation of, meta3C and Hi-C data. We provide a strategy to improve the signal-to-noise ratio by discarding reads that map to repetitive elements and to the end of contigs. We also show the importance of using spike-in controls to quantify whether the cross-linking step in meta3C and Hi-C protocols has been successful.

After filtering for spurious links, 87 ARGs were linked to their bacterial hosts across all datasets, including 27 ARGs in the meta3C dataset we generated. We show that commensal gut bacteria are an important reservoir for ARGs, with genes encoding for aminoglycoside and tetracycline resistance being widespread in anaerobic commensals of the human gut.

## Introduction

The gut microbiota is a complex ecosystem that is frequently characterised through high-throughput shotgun sequencing to quantify and characterise the abundance of viruses, bacteria, fungi, and protists [1]. Using sequencing-based approaches, it remains a challenge to link genes in the gut microbiota to their microbial hosts as metagenomic assemblies are often highly fragmented and metagenome assembled genomes (MAGs) are frequently incomplete or suffer from contaminating sequences [2,3]. While contiguity of assemblies can be improved by the incorporation of long-read sequencing data [4,5], the hosts of plasmids can only be predicted, but not conclusively identified, by a variety of bioinformatic approaches [6]. The linkage of genes to their microbial hosts is particularly important for genes that confer antibiotic-resistant phenotypes to their hosts. Sequencing-based studies have shown that the human gut microbiota forms a reservoir of antibiotic resistance genes (ARGs) [7,8]. These genes often have the potential to spread promiscuously in microbial populations, particularly when they are associated with plasmids [9]. Horizontal transfer of antibiotic resistance genes in the gut has been observed among Enterobacteriaceae, *Bacteroides* and *Enterococcus* strains [10–12]. It is thus of interest to disentangle ARG-host linkage across the gut microbiota with the long-term goal to understand to what extent commensal bacteria can serve as a conduit for ARGs to be transferred to gut-dwelling opportunistic pathogens like *Clostridioides difficile, Escherichia coli, Klebsiella pneumoniae, Enterococcus faecalis* and others [2].

To improve linkage of genes to their hosts in microbial ecosystems, metagenomic proximity ligation techniques have been developed [13]. In short, these techniques cross-link DNA within microbial cells through an incubation with formaldehyde, followed by digestion with restriction enzymes, proximity ligation, the reversal of crosslinks by treatment with a protease and finally high-throughput sequencing of the resulting fragments (Figure 1). Metagenomic chromosome confirmation capture was first described by Marbouty and colleagues and was termed meta3C [14]. Shortly afterwards, other protocols were published that include an additional step to specifically enrich for cross-linked DNA, which involves tagging the termini of DNA fragments with biotin before ligation [15,16]. After removal of the cross-links, DNA is sheared and streptavidin beads are used to pull down biotin-tagged fragments, thus enriching for ligation junctions. Protocols with this additional enrichment step are collectively termed Hi-C.

**Figure 1.**
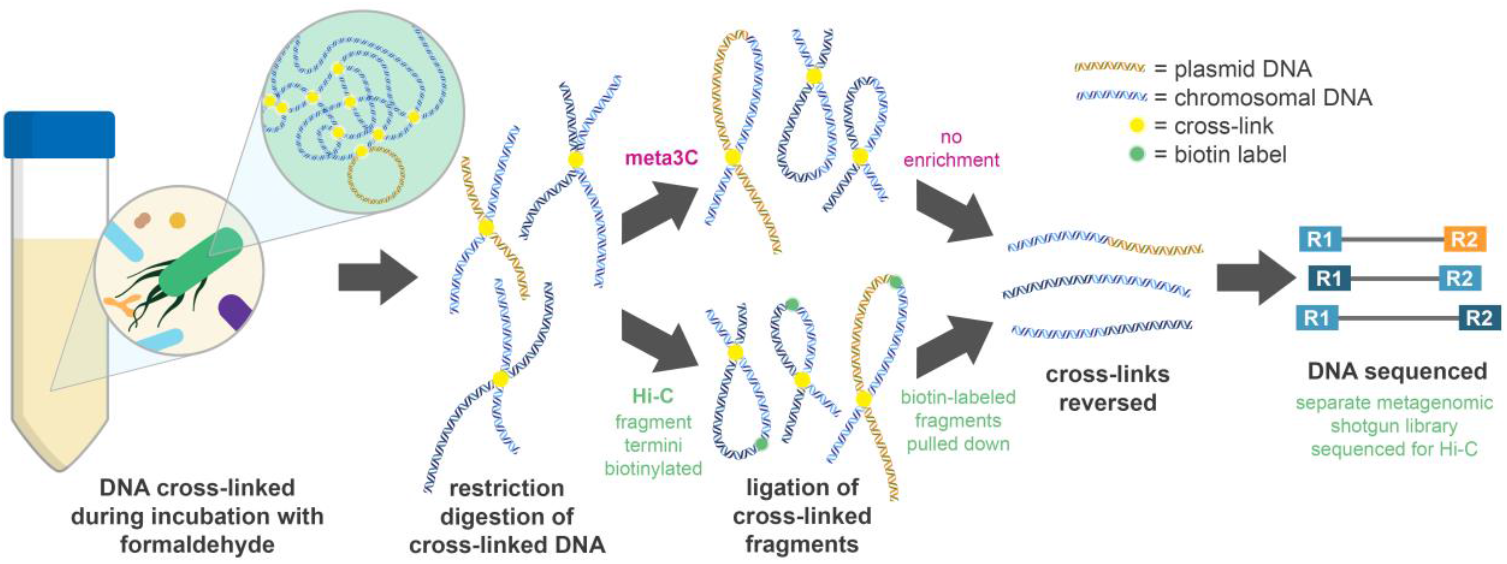
Metagenomic chromosome conformation capture approaches. Formaldehyde is used to cross-link DNA-bound proteins before cell lysis and enzymatic digestion of the DNA. In meta3C, the cross-linked digested fragments are then ligated. In Hi-C, the digested fragments are tagged with biotin prior to ligation, enabling enrichment of ligated biotin-labelled fragments following ligation and DNA shearing. The cross-links are then removed during treatment with a protease, and the fragments undergo high-throughput sequencing.

Due to its lower cross-linking efficiency a meta3C library must be sequenced more deeply than a Hi-C library to ensure that sufficient numbers of cross-linked fragments are sequenced [13]. However, the relatively low proportion of non-cross-linked fragments sequenced from a meta3C library allows generation and scaffolding of contigs directly from meta3C sequencing data [14]. For Hi-C, additional shotgun sequencing of the sample is required for metagenomic assembly, which must then be analysed in conjunction with the Hi-C data to link assembled contigs [15,16]. 3C/Hi-C approaches have been used to considerable effect in improving the assembly of MAGs in complex microbial ecosystems such as those present in the bovine rumen [17], the gut of dogs [18], sheep [19] and pigs [20]. Several studies have been performed using 3C/Hi-C to study the human gut microbiota [21–25].

The overarching goal of this study was to explore whether meta3C and Hi-C can be used to reliably link ARGs to their microbial hosts. To this aim, we generated a meta3C dataset, using spike-in controls, of a human stool sample. We combined the analysis of this dataset with re-analysis of publicly available meta3C and Hi-C data generated using gut microbiome samples, to identify and address technical challenges in the analysis of metagenomic chromosome conformation capture data to determine linkage between ARGs and chromosomes and plasmids of their microbial hosts.

## Results

### Generation of a meta3C library using a human stool sample

A meta3C library was generated using a human stool sample from an individual suffering from Inflammatory Bowel Disease (IBD). Prior to the first step of the meta3C protocol (i.e. incubation with formaldehyde), we spiked in two antibiotic resistance gene carrying strains, *E. coli* E3070 [26] and *E. faecium* E745 [27], at 6.4 × 10^8^ colony forming units/g each, equivalent to an estimated 0.5% of the total community. Two meta3C libraries, differing by the enzymes (MluCI and HpaII) used for restriction digestion were generated and sequenced independently. After processing of the reads (to remove low-quality, duplicate, and human reads), 101 million and 97 million high-quality reads remained for the HpaII and MluCI meta3C libraries, respectively. These reads were then combined to generate the G_3C dataset and used for the metagenomic assembly. The reads were assembled into 89,005 contigs, with a contig N50 of 10,778 and a total length of 404,824,063 bp (Table 1).

**Table 1.** Read counts and assembly statistics.

| Dataset | Assembly | Read length (bp) | Raw reads | Processed reads | Total length of assembly (bp) | No. of contigs | Contig N50 | Reference |
| --- | --- | --- | --- | --- | --- | --- | --- | --- |
| G_3C | **G_3C** | 2 × 150 | 223,169,682 | 198,493,086 | 404,824,063 | 89,005 | 10,778 | This study |
| M_3C | **M_3C** | 2 × 75 | 375,815,400 | 366,961,002 | 480,933,195 | 116,057 | 7,562 | [28] |
| P_HiC | P_HiC | 2 × 150 | 171,853,886 | 157,755,162 | ND | ND | ND | [24] |
|  | **P_SG** | 2 × 150 | 250,884,672 | 237,293,522 | 528,999,126 | 104,368 | 14,455 |  |
| Y_3C | **Y_3C_A** | 2 × 160 | 3,019,738,680 | 2,921,579,828 | 1,040,533,919 | 177,689 | 17,376 | [21] |
|  | Y_SG_A | 2 × 160 | 416,571,650 | 410,280,634 | 658,813,968 | 101,575 | 22,551 |  |
|  | **Y_3C_B** | 2 × 160 | 682,773,219 | 1,239,950,680 | 866,666,497 | 146,989 | 18,851 | [21] |
|  | Y_SG_B | 2 × 160 | 202,617,904 | 198,775,312 | 484,955,068 | 83,017 | 19,100 |  |
| D_HiC | D_HiC | 2 × 80 | 143,286,468 | 133,509,800 | ND | ND | ND | [23] |
|  | **D_SG** | 2 × 150 | 20,088,550 | 18,925,950 | 131,298,239 | 37,723 | 5,924 |  |
| K_HiC<br>(average*) | K_HiC | 2 × 150 | 41,021,508 | 37,984,239 | ND | ND | ND | [22] |
|  | **K_SG** | 2 × 150 | 90,510,991 | 83,397,427 | 156,853,335 | 32,197 | 18,026 |  |

\*for K\_HiC, an average of 43 samples is presented in this table; x\_SG = accompanying shotgun metagenomic reads; assemblies in bold were used during analysis of 3C/Hi-C data; bp = base pairs; ND = not determined

### 3C/Hi-C datasets reflect composition of the gut microbiota

To benchmark the meta3C data generated here against previously published 3C/Hi-C gut microbiota data, several published datasets using gut microbiota samples that were available at the inception of this study (June 2020) were reanalysed (Table 1). All these datasets originated from humans, with the exception of M_3C, which was generated using a murine gut microbiota sample [28].

The raw reads from these published datasets were downloaded from NCBI, processed and analysed identically to the reads generated in this study. Where more than one enzyme was used and sequenced as separate libraries, or if 3C/Hi-C datasets were made up of technical repeats, the reads were combined before metagenomic assembly. For studies using the meta3C protocol, assemblies were made using the meta3C data, as advised in the publication describing meta3C [14], while for Hi-C data assemblies were generated using the shotgun metagenomic sequencing datasets (Table 1). The study of Yaffe and Relman [21] was the only study which contained both meta3C and shotgun sequencing data. We assembled both but we decided to use the assembly based on meta3C data for further analyses as these assemblies were 1.6-fold and 1.8-fold larger (for sample A and B, respectively) than the assemblies generated by shotgun sequencing data.

The taxonomic compositions of the gut microbiota, on the basis of the processed reads from all datasets, were determined using MetaPhlAn3 [29]. Among classified reads, most samples showed results that can be expected for a human faecal sample, with the majority of the reads being assigned to the Clostridia and Bacteroidia classes (Figure 2). Some samples differed greatly from the others, such as K_HiC_N1-4, where 88.55% of the classified reads were assigned to ‘Viruses_unclassified’, which may reflect the neutropenic nature of most of the individuals in the K_HiC dataset [22]. For the dataset generated for this study (G_3C), 39.92% of classified reads were assigned to the class Clostridia, 17.63% to Actinobacteria, 10.64% Coriobacteria, and 9.94% to Bacteroidia (Figure 2). The *Enterococcus* and *Escherichia* genera had similar abundances to each other (3.79% and 3.43%, respectively), which suggests that the *E. coli* and *E. faecium* strains had been spiked in at a higher level than the 0.5% target due to an overestimation of the total bacterial loads.

**Figure 2.**
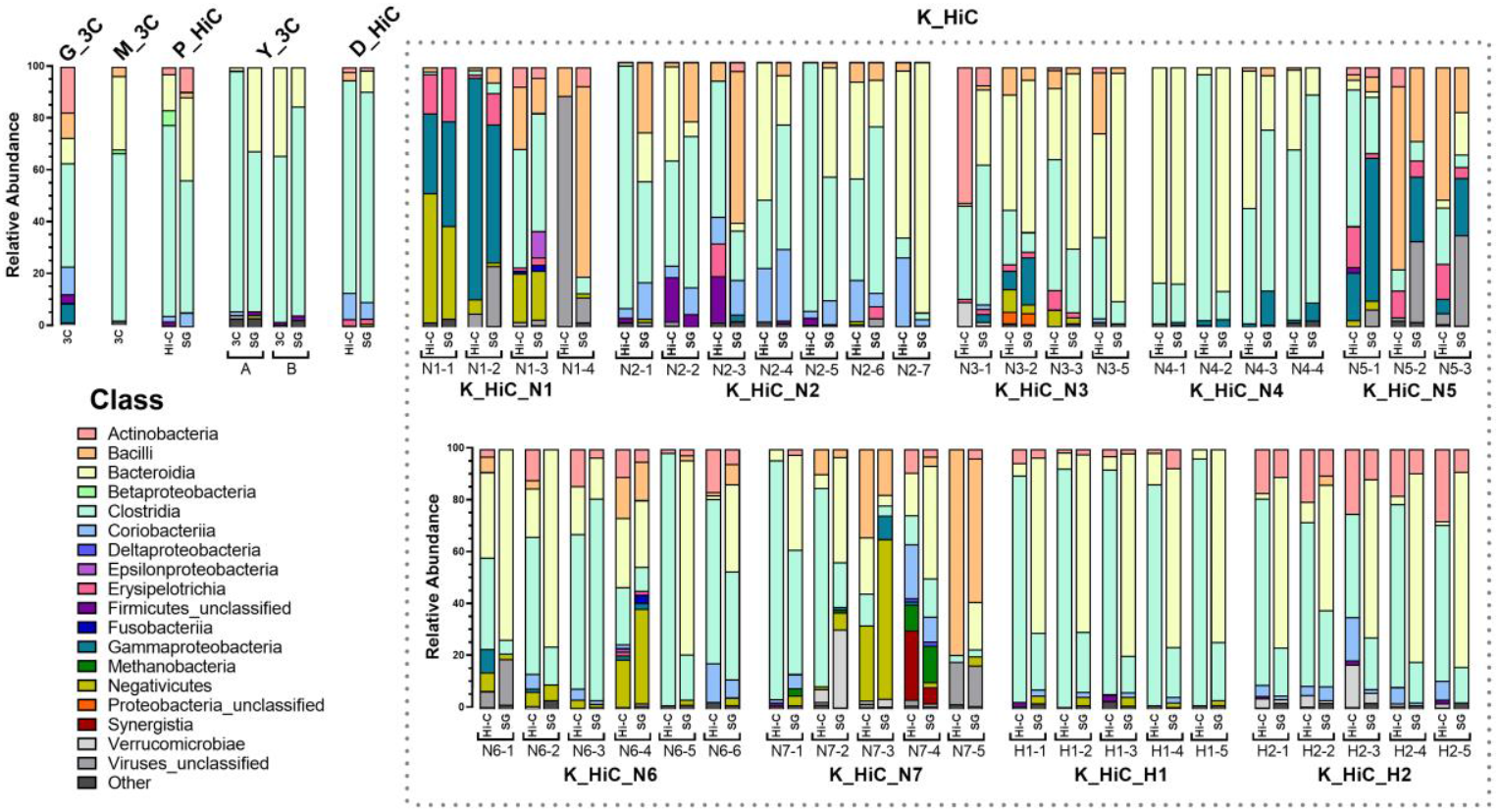
Class-level compositions of all datasets. The reads from all datasets were taxonomically profiled using MetaPhlAn3. The stacked bars show the relative abundance (%) of each class for the classified reads. Reads that could not be classified by MetaPhlAn3 (∼60% of reads for each dataset) are excluded. For the K_HiC dataset, individuals are either neutropenic (N1-7) or healthy (H1-2) with multiple samples collected longitudinally for each individual.

### Diverse antibiotic resistance genes are present in all datasets

After phylogenetic profiling of the reads, ABRicate was used to identify contigs containing ARGs in the metagenomic assemblies. In the G_3C assembly, 37 contigs containing ARGs were identified. The known ARGs from the E3090 and E745 spike-ins were all present (Figure 3). For E745, the two chromosomal ARGs (*aac(6’)-Ii* and *msr*(C)) had similar abundances of 15.0 and 14.3 reads per kilobase per million mapped reads (RPKM), respectively. The other ARGs from E745, *vanHAX* and *dfrG*, are carried on plasmids, and had higher abundance (43.4 and 34.9 RPKM) than the chromosomal ARGs, likely due to being carried on a plasmid that has a higher copy number than the chromosome. For the E3090 ARGs, six chromosomal ARGs (*sul1, sul2, ant(3’’)-Ia, bla*_OXA-1_, *floR*, and *mdf*(A)) had relatively similar abundances, ranging from 9.6-27.4 RPKM. The ARGs carried on plasmids in E3090 had higher relative abundances. The *mcr-1*.*1* gene had an abundance of 98.1 RPKM, while *bla*_TEM_ was present at a high abundance of 104.8 RPKM. The *bla*_TEM_ gene present in the metagenomic assembly was identified as *bla*_TEM-116_, as opposed to *bla*_TEM-1B-_ that the E3090 genome contains. These genes differ by 5 single nucleotide polymorphisms (SNPs), so this is most likely due to a misassembly in either the original genome sequence or the metagenomic assembly.

**Figure 3.**
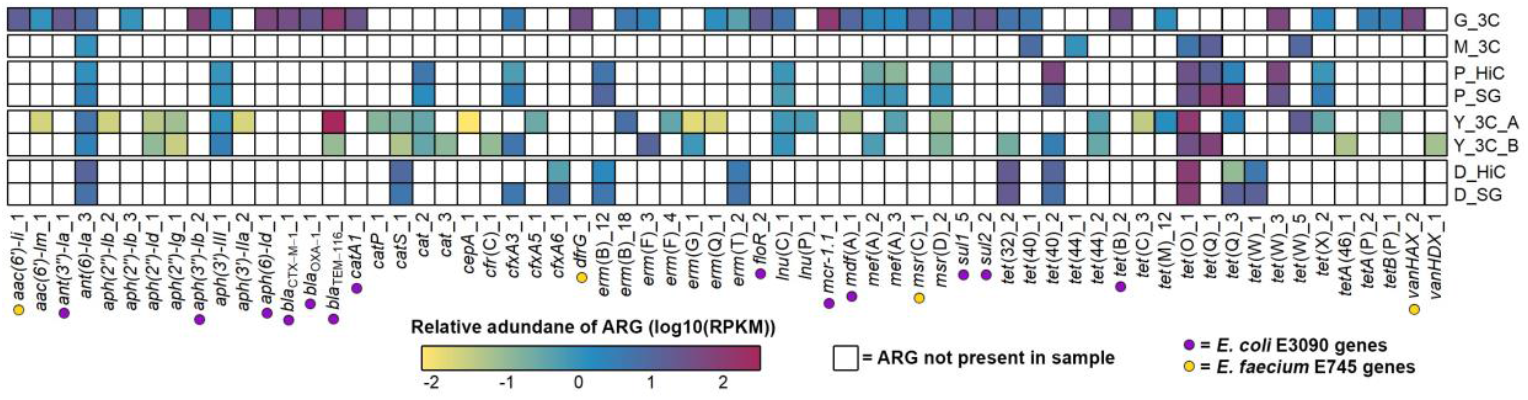
Relative abundance of antimicrobial resistance genes (ARGs) in 3C/Hi-C datasets. The ARG sequences from the assemblies of each dataset were isolated, and the reads from that dataset were mapped to the ARGs (columns). The relative abundance was calculated as reads per kilobase per million mapped reads (RPKM). White cells mean the ARG was not present, and coloured cells show that the ARG was present, with the colour relating to the relative abundance of the ARG within that set of reads (log(10) transformed RPKM values). Different datasets are separated by gaps in the heatmap. 3C datasets (*_3C) have rows showing the RPKM of the 3C reads mapping to the ARGs identified in the 3C metagenomic assembly. Hi-C datasets show RPKM of the shotgun reads (*_SG) or Hi-C reads (*_HiC) mapping to ARGs identified in the shotgun metagenomic assembly. The ARGs highlighted with a coloured dot are ARGs from the spike-ins in the G_3C dataset (purple = *E. coli* E3090, yellow = *E. faecium* E745).

The rest of the datasets contained many and diverse ARGs, with 71 unique ARGs in total across the datasets, excluding the K_HiC samples (Figure 3). The 86 samples in the K_HiC dataset (43 Hi-C and 43 corresponding shotgun metagenomic samples) contained 141 unique ARGs and have been shown separately in Supplementary Figure 1.

### Presence of spurious crosslinks in 3C/Hi-C data

To identify reads from cross-linked fragments of DNA, the first 50 bp of the 3C/Hi-C reads from each dataset were first mapped against their respective metagenomic assemblies. For Hi-C datasets (P_HiC, D_HiC, K_HiC), the Hi-C reads were mapped to assemblies generated from the accompanying shotgun metagenomic library, whereas 3C reads from the 3C datasets (G_3C, M_3C, Y_3C) were mapped to assemblies generated directly from the 3C library. From the reads that mapped with a mapping quality (MAPQ) >20, intercontig read pairs were identified as instances where both reads of the pair mapped to different contigs (Figure 4; Supplementary Table 1), indicating the read pair potentially came from a cross-linked fragment of DNA.

**Figure 4.**
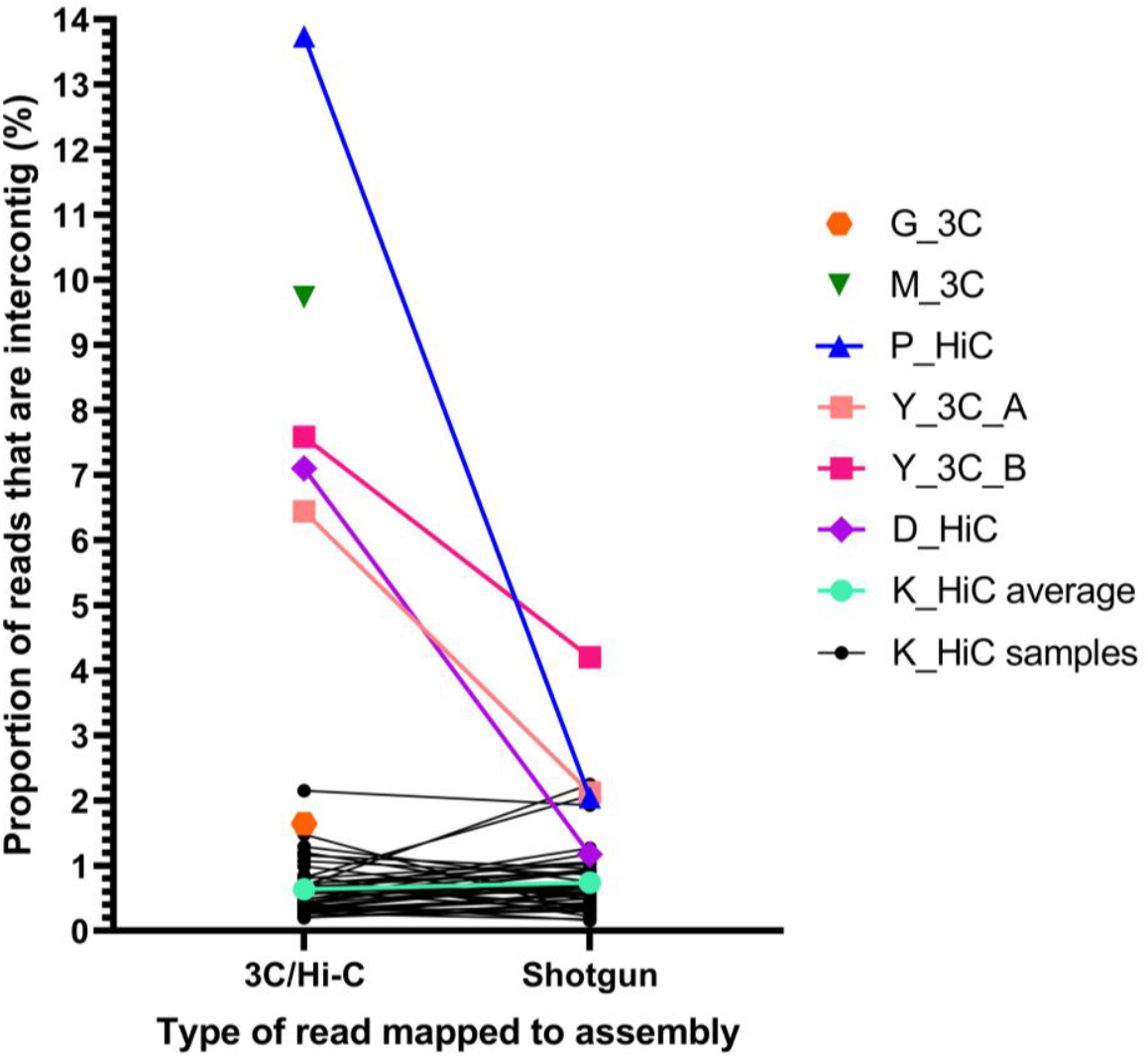
Proportion of intercontig reads in 3C/Hi-C and shotgun reads of the same sample. The first 50 bp of each read were mapped against the corresponding assembly, and pairs where each read of the pair mapped to different contigs were labelled as intercontig reads. Y-axis shows the percentage reads that were intercontig. K_HiC average (cyan) is the average for all 43 K_HiC samples (black). G_3C (orange) and M_3C (green) did not have accompanying shotgun reads, so only the intercontig proportion for the 3C reads are shown.

The proportion of intercontig reads greatly varied across the datasets, with the highest being 13.74% for P_HiC, and the lowest being 0.2% for K_HiC_N1-1 (0.64% average across all K_HiC samples). For the meta3C datasets, M_3C had the highest proportion of intercontig reads at 9.73%. The G_3C dataset had the lowest number of cross-linked reads of the meta3C datasets at 1.65% (Supplementary Table 1).

Due to the large differences in the proportion of intercontig reads across the datasets, we set out to study whether these intercontig reads were truly a result of physical cross linking. We first mapped shotgun metagenomic reads, which, by definition, cannot have been physically cross-linked, in the datasets that contained them (Y_3C_A/B, D_HiC, P_HiC, K_HiC) back to the assemblies in the same way as the 3C/Hi-C reads were in the previous step. They were then analysed the same as the 3C/Hi-C reads to isolate the intercontig read pairs and calculate the proportion of intercontig reads. The shotgun metagenomic reads showed a background level of 0.16 to 4.20% intercontig reads (Figure 4). These reads have not been generated from physically cross-linked fragments of DNA and shall thus be referred to as ‘spurious intercontig reads’. In the K_HiC datasets, the average proportion of intercontig reads from the shotgun metagenomic reads was 0.74%, compared to the average of 0.64% cross-linked reads from the Hi-C reads. This suggested that there may be no, or very few, reads resulting from the physical cross-linking of DNA in the K_HiC dataset.

### Non-contiguous assemblies introduce noise in 3C/Hi-C datasets

We recognised that the G_3C dataset, with an intercontig read proportion of 1.65%, is within the range of the spurious intercontig reads from the shotgun metagenomic data and may thus also have been insufficiently cross-linked during the experimental procedure. Because the G_3C dataset contained the *E. coli* and *E. faecium* spike-ins, for which whole genome sequences are available, the intercontig reads mapping to the respective genome sequences could be examined further. G_3C reads were mapped to the *E. coli* E3090 and *E. faecium* E745 genomes to isolate spike-in 3C reads for each genome. These reads were then compared to whole genome sequencing (WGS) reads downloaded from NCBI for each genome by mapping the first 50 bp of all reads to the G_3C assembly (Supplementary Table 2).

The proportion of intercontig reads from the 3C reads (0.98% and 0.99% for *E. coli* and *E. faecium*, respectively) were comparable to the WGS reads (0.78% and 2.34% for *E. coli* and *E. faecium*, respectively) confirming again that short-read sequencing produces a considerable background level of reads that can be erroneously interpreted as originating from cross-links. Aligning both the intercontig and non-intercontig reads from the G_3C spike-in and the WGS reads back to their respective genomes revealed the regions the reads were mapping to on the genome. Both the intercontig and non-intercontig reads spanned the whole genome for both spike-ins and aligned to different genomic regions (Figure 5). A greater proportion of the intercontig reads mapped to insertion sequence elements (IS elements) in the genome compared to the non-intercontig reads for all sets of reads, except for the G_3C E3090 reads. This was most clear in the E745 reads, where over 20% of the intercontig reads for both the G_3C and WGS reads aligned to IS elements, compared to less than 1% of the non-intercontig reads, suggesting that the presence of multiple copies of IS elements in the assembly is partially responsible for the spurious intercontig reads. Using ABRicate with the ISfinder database [30], 93 copies of IS elements (18 different types) were found across the 3,168,411-bp *E. faecium* E745 genome (29.3 IS elements/Mbp), compared to 79 IS element copies (25 different types) in the 5,270,976 bp *E. coli* E3090 genome (15.0 IS elements per Mb). The E745 reads thus have a higher chance of mapping to an IS element, causing more spurious intercontig reads.

**Figure 5.**
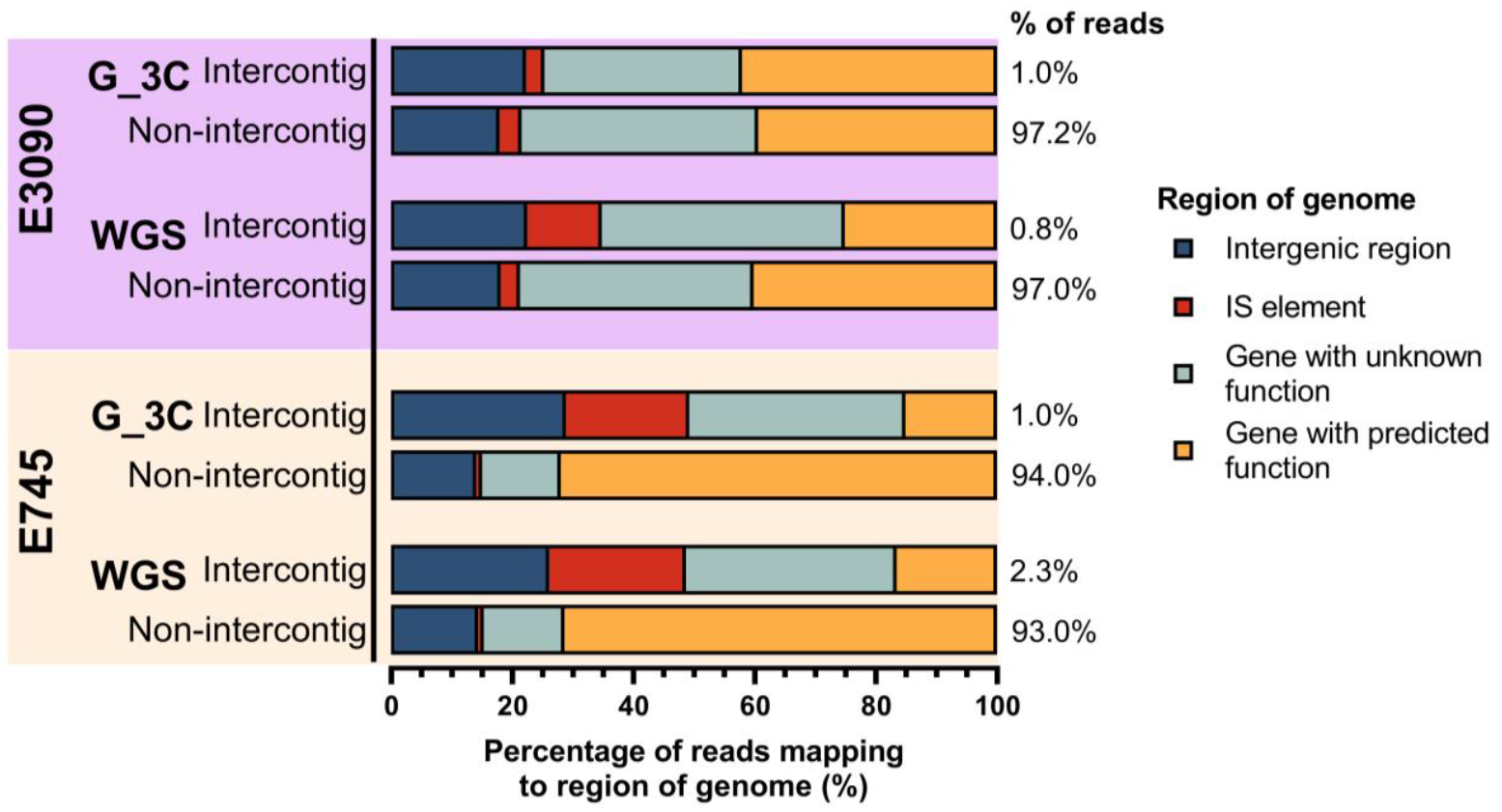
G_3C reads and whole genome sequencing reads mapping to genome sequences of spike-in controls. Both the intercontig and non-intercontig reads for G_3C spike-in reads and WGS reads of the spike-ins were mapped to their respective genomes. The genomes were annotated using Prokka and the regions in which the reads mapped to were grouped into four categories (see legend). IS element = Insertion Sequence element. Percentages at the end of the stacked charts show the proportion of total reads that were assigned to intercontig/non-intercontig.

Next, the position in the contigs from the G_3C assembly that the spike-in reads mapped to was checked to determine whether spurious intercontig reads were more likely to map near to the beginning or end of a contig, meaning they were potentially caused by fragmentation in the assembly. Indeed, a greater proportion of the intercontig reads for both the G_3C and WGS spike-in reads mapped within 500 nt of the ends of a contig compared to the non-intercontig reads (Figure 6). For G_3C E3090 reads, 37.7% of the intercontig reads mapped within the first or last 500 nt of a contig, compared to only 5.0% of non-intercontig G_3C E3090 reads. This observation was even clearer for the E3090 WGS and G_3C/WGS E745 reads, where over 80% of the intercontig reads were mapping near the ends of a contig, compared to less than 10% of the non-intercontig reads (Figure 6).

**Figure 6.**
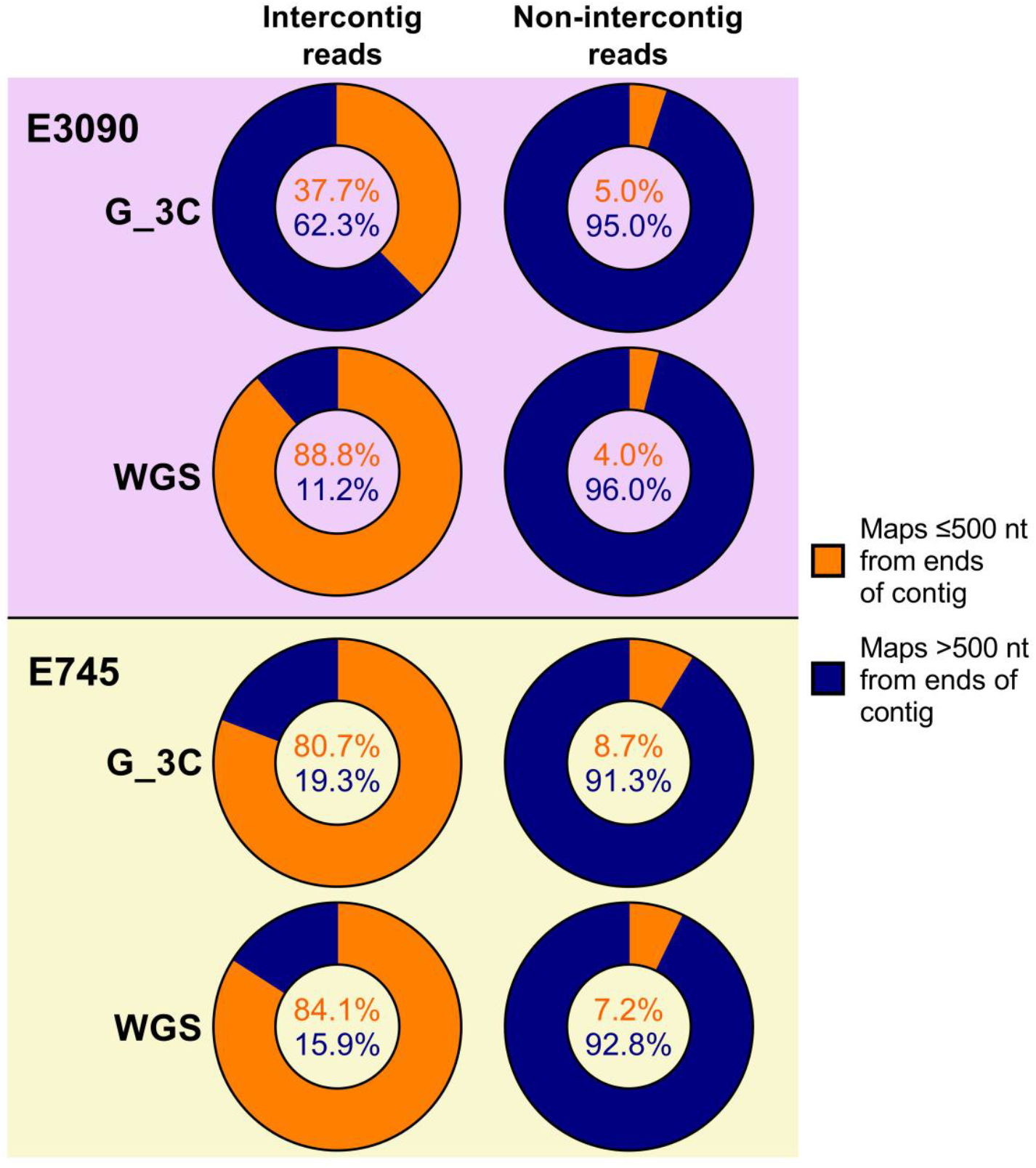
Proportions of reads mapping within the first or last 500 nucleotides (nt) of a contig in the G_3C assembly for spike-in G_3C and whole genome sequencing (WGS) reads. The position of the alignment to contigs in the G_3C assembly was checked for both intercontig and non-intercontig read pairs from WGS reads and reads from G_3C that mapped to each spike-in genome. Orange shows the proportion of reads mapping within 500 nt of the ends of a contig. Blue shows the proportion of reads mapping more than 500 nt away from the ends of a contig.

To determine whether intercontig reads mapped to the ends of contigs for all 3C/Hi-C reads, the positions in the metagenomic assembly that the reads mapped to were checked for all datasets. The majority of spurious intercontig reads from the shotgun metagenomic data mapped within the first or last 500 nt of a contig for all datasets that had shotgun data (Figure 7). For the 3C/Hi-C intercontig reads, the proportion varied, but was lower for P_HiC, D_HiC, Y_3C_B, and M_3C (12.7%, 27.5%, 32.1%, and 19.4%, respectively), compared to around 52% for G_3C and Y_3C_A.

**Figure 7.**
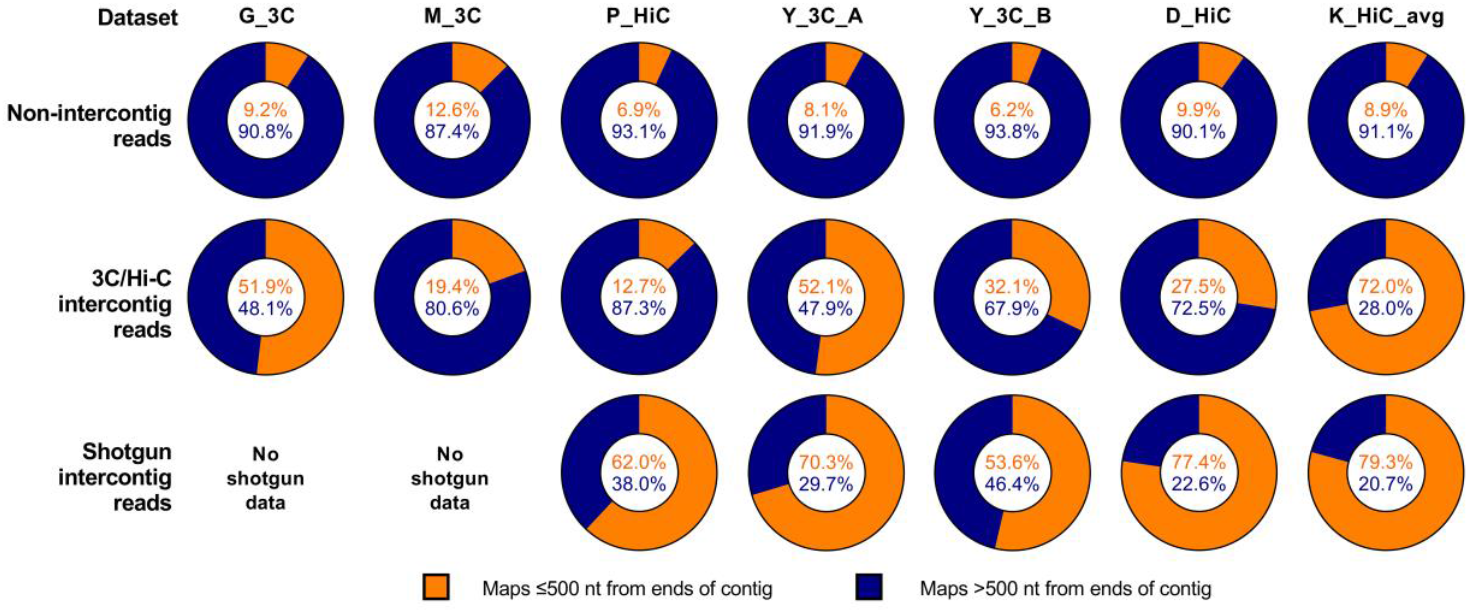
Proportions of intercontig reads mapping within the first or last 500 nucleotides (nt) of a contig in their respective assemblies for all datasets. The position of the alignment to contigs was checked for the intercontig reads in all datasets. Orange shows the proportion of reads mapping within 500 nt of the ends of a contig. Blue shows the proportion of reads mapping greater than 500 nt away from the ends of a contig.

The proportion of intercontig 3C/Hi-C reads mapping near ends of a contig correlated (*R*^2^ = 0.86; P = 0.0028) with the proportion of intercontig reads in the dataset, whereby datasets with a higher proportion of intercontig reads in the sequence data had a lower proportion of intercontig reads mapping near the ends of a contig (Supplementary Figure 2). This, along with the high proportion of shotgun intercontig reads mapping near the ends of a contig, suggested that many spurious intercontig reads can be filtered out by removing those that mapped within the first or last 500 nt of a contig.

### Filtering reads that map in the first 500 nt of a contig removes most spurious intercontig reads

To reduce the number of spurious intercontig reads in the data, intercontig reads that mapped within the first 500 nt of a contig were removed in all datasets, reducing the proportion of intercontig reads by 32.6%, on average, across all the datasets (Figure 8). Notably, when the same filtering step was performed on the shotgun data, the proportion of intercontig reads decreased by 63.8%, suggesting that this step is essential to reduce the number of spurious intercontig reads in the data. After removing the reads mapping near the ends of contigs, the proportion of intercontig reads from the Hi-C data in the K_HiC dataset was 0.18% on average, hardly different from the average of the K_SG spurious intercontig reads (0.16**%**). Therefore, this dataset was not included in further analyses.

**Figure 8.**
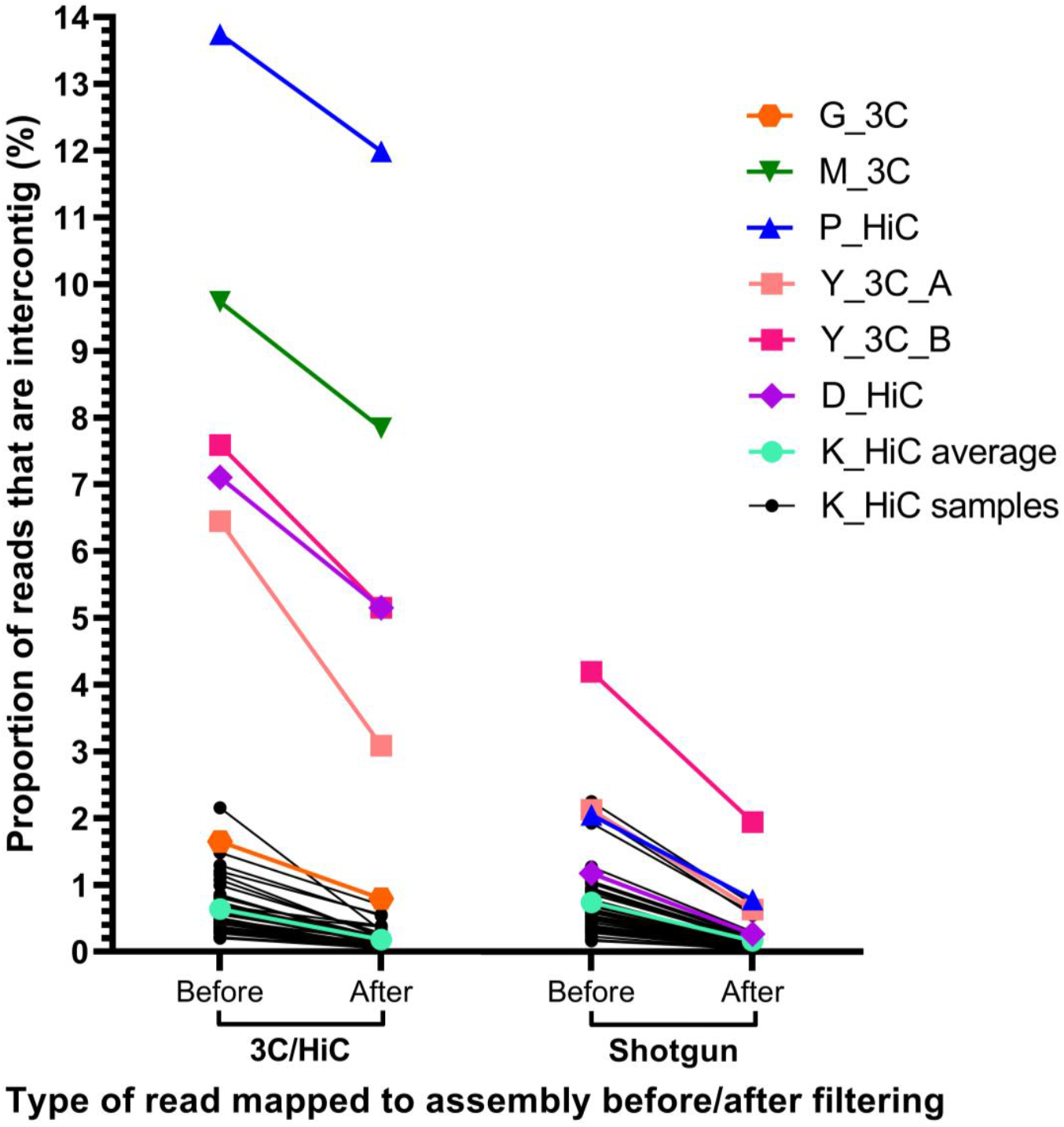
Proportion of intercontig reads in 3C/Hi-C and shotgun reads before and after filtering. The first 50 bp of each read were mapped against the corresponding assembly, and pairs where each read of the pair mapped to different contigs were labelled as intercontig reads (‘Before’ on X-axis). These were then filtered to remove intercontig reads that mapped within the first or last 500 nt of a contig (‘After’ on X-axis). Y-axis shows the percentage reads that were intercontig. K_HiC average (cyan) is the average for all 43 K_HiC samples (black). G_3C (orange) and M_3C (green) did not have accompanying shotgun reads, so only the intercontig proportion for the 3C reads before and after filtering are shown.

### Linking ARGs to microbial hosts in 3C/Hi-C datasets

After filtering out the intercontig reads that mapped to the 500 nt ends of contigs, pairs where one read mapped to an ARG contig in its respective assembly were identified. To further reduce the impact of the noise from any remaining spurious intercontig reads, contigs were only considered linked to ARG contigs if there were ≥5 unique intercontig read pairs linking them. In addition, ARG-linked contigs identified as IS elements were also filtered out (Table 2).

**Table 2.** Number of contigs linking to ARG contigs in 3C/Hi-C datasets.

| Dataset | G_3C | M_3C | P_HiC | Y_3C |  | D_HiC |
| --- | --- | --- | --- | --- | --- | --- |
|  |  |  |  | Y_3C_A | Y_3C_B |  |
| Intercontig reads linking contigs to an ARG contig | 26,607 | 17,321 | 28,200 | 186,177 | 128,774 | 19,475 |
| Unique contigs linked to ARG contig | 4,767 | 6,763 | 4,517 | 9,172 | 14,757 | 3,007 |
| Linked $\geq 5$ times | 519 | 264 | 445 | 611 | 1,661 | 392 |
| After removal of links to IS elements | 466 | 264 | 443 | 606 | 1,655 | 392 |
| After removal of links to plasmids | 342 | 259 | 439 | 594 | 1,627 | 387 |
| Number of ARGs linked to host(s) ( / number of ARGs in sample) | 27 / 37 | 6 / 7 | 9 / 15 | 23 / 30 | 16 / 23 | 6 / 11 |

For datasets that used multiple restriction enzymes, numbers presented are a combined total; ARG = antimicrobial resistance gene; IS element = insertion sequence element

**Table 3.** Published 3C datasets downloaded in study.

| Study | Accession number | Reference |
| --- | --- | --- |
| M_3C | PRJNA302158 | [28] |
| P_HiC | PRJNA413092 | [24] |
| Y_3C | PRJNA505354 | [21] |
| D_HiC | PRJNA377403 | [23] |
| K_HiC | PRJNA649316 | [22] |

For G_3C, this resulted in 26,607 intercontig reads that linked a total of 466 contigs to 27 out of 37 of the ARG contigs (Table 2). Linked contigs that mapped with >99% identity to known plasmid sequences in the NCBI nt database, which were all linking to ARGs from the spike-ins, were removed as no definitive identification of the microbial hosts could be made (Table 2). The remaining contigs were then taxonomically classified using Kraken2. This revealed that the ARGs were linked to a wide range of taxa (Figure 9). Genes from the E745 spike-in were correctly linked to *Enterococcus*, although *vanHAX* was excluded as it only linked to plasmid contigs. The same was true for *catA1* and *bla*_TEM_ in the *E. coli* E3090 spike-in, however the remaining E3090 ARGs were all linked to *Escherichia*. A small proportion (1.7-3.7%) of the contigs that linked to several of the E3090 ARGs (*bla*_CTX-M-1_, *mcr-1*.*1, aph(3’’)-Ib, aph(6)-Id, mdf*(A), *sul1, ant(3’’)-Ia, bla*_OXA-1_, and *sul2*) were only classified to the family level as Enterobacteriaceae, with the remaining contigs linked to these genes being successfully classified to species level as *E. coli*.

**Figure 9.**
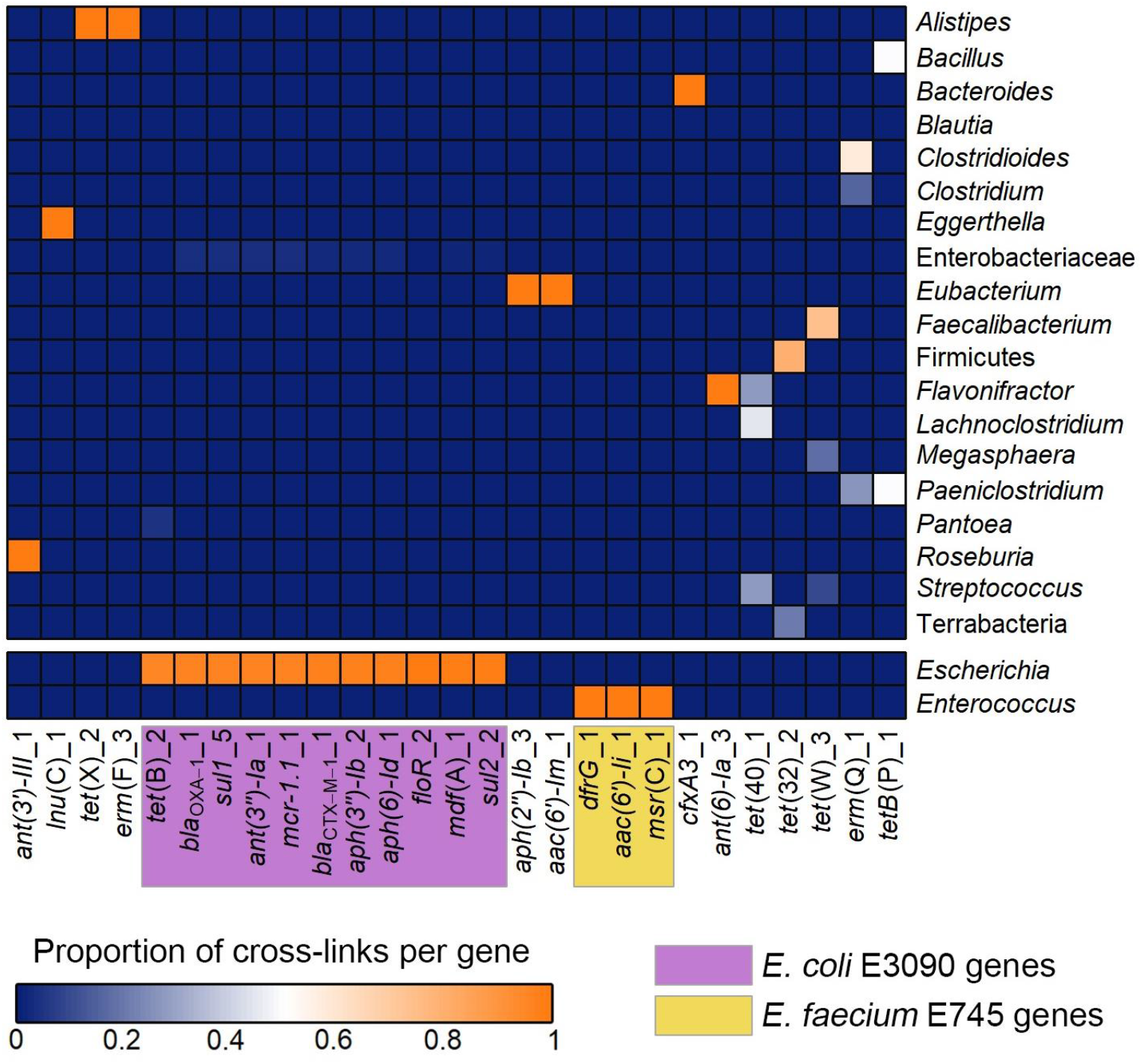
Heatmap showing antimicrobial resistance genes (ARGs) linked with their microbial hosts for G_3C. Contigs linked to ARG-containing contigs were taxonomically classified using Kraken2. The heatmap shows the proportion of contigs linked to each ARG that was classified as the taxon on the right. *Enterococcus facecium* E745 and *Escherichia coli* E3090 were spiked into the stool sample, and the ARGs that these strains carried are highlighted in yellow and purple, respectively.

These results indicated that the analysis pipeline used here could successfully link the spike-in ARGs to their correct host. The non-spike-in ARGs linked to a wide range of hosts. Some ARGs such as *cfxA3* and *tet*(X) linked to single hosts, whereas others, like *tet*(40) and *tet*(W), were widespread and linked to various gut commensals. Where ARGs were associated with multiple taxa, the potential microbial hosts were usually related at phylum level, such as *tet*(40) which linked to the genera *Streptococcus, Flavonifractor*, and *Lachnoclostridium*, which are all in the Firmicutes phylum.

ARGs were then linked to their microbial hosts for the other 3C/Hi-C datasets. As with G_3C, some ARGs were linked to few microbial hosts, whereas others were linked to a wide range of hosts (Figure 10), and the proportions of ARGs successfully linked to their hosts were high, with 6/11, 9/15, 23/30, 16/23, and 6/7 for D_HiC, P_HiC, Y_3C_A, Y_3C_B, and M_3C, respectively (Table 2).

**Figure 10.**
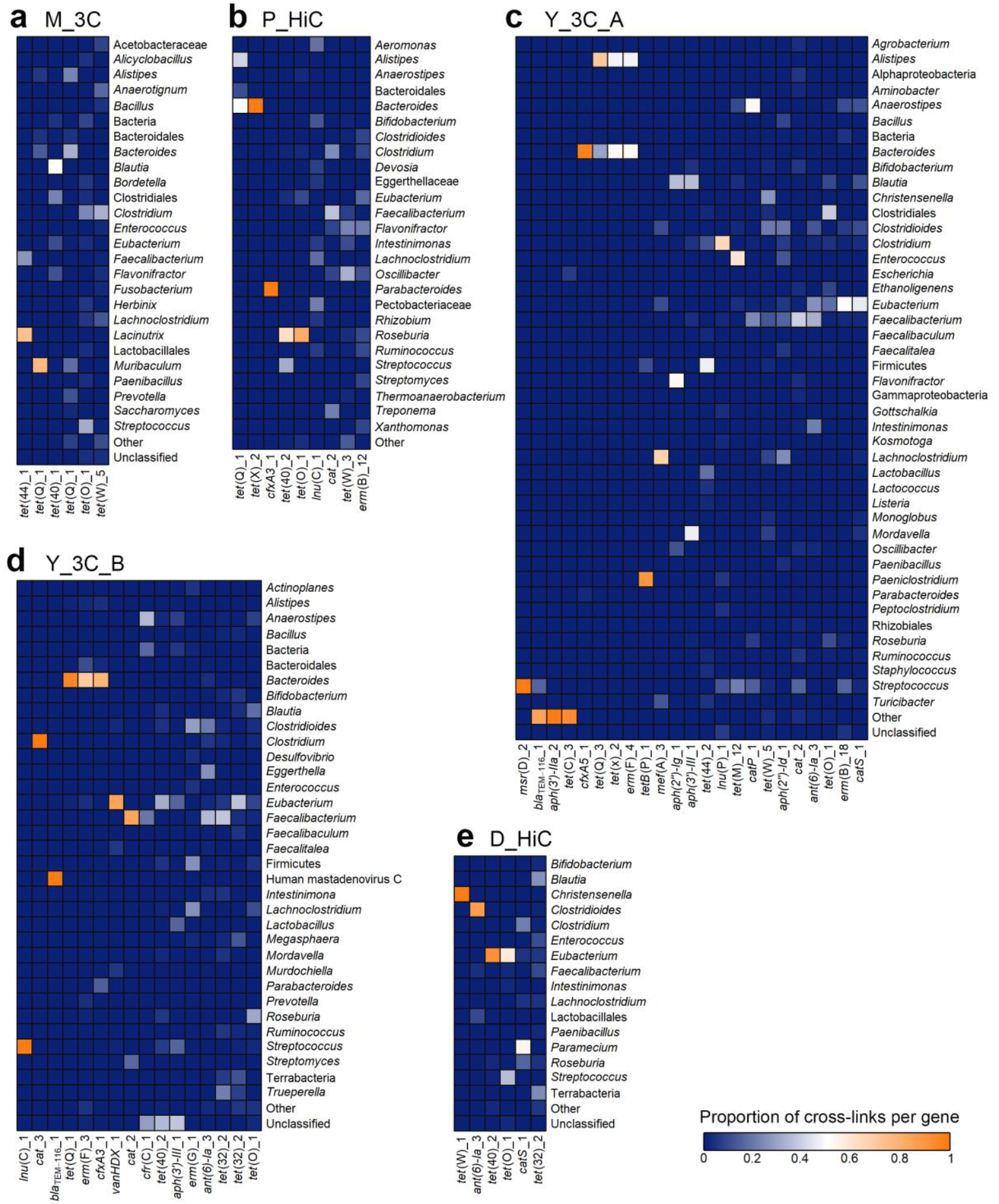
Heatmap showing antimicrobial resistance genes (ARGs) linked with their microbial hosts for downloaded 3C/Hi C datasets. Contigs linked to ARG-containing contigs were taxonomically classified using Kraken2. The heatmaps show the proportion of contigs linked to each ARG that was classified as the taxon on the right. Where there were multiple taxa that made up a proportion of no more than 0.02 for any ARG in that dataset, they have been grouped into “Other”.

Some of the shared ARGs linked to the same hosts across datasets, whereas others linked to multiple diverse hosts. The *tet*(X), *tet*(Q), and *erm*(F) genes were linked predominantly to *Alistipes*, and *Bacteroides*, both from the order Bacteroidales, in all datasets that they were present in. The beta-lactamase *cfxA3* was only linked to *Bacteroides* in all datasets that it was present in. Conversely, *tet*(O), *tet*(40), *lnu*(C), *cat, ant(6)-Ia*, and *tet*(W) showed a wide range of hosts across the datasets, with *tet*(W) linking to over 20 taxonomic classifications in total across five datasets.

Overall, these results indicate that the ARGs identified in the assemblies were able to be linked to their microbial hosts using meta3C/Hi-C data, with stringent filtering to minimise the impact of spurious links, revealing some genes to be promiscuous and linking to a wide range of gut bacteria.

## Discussion

Previous studies have implemented 3C/Hi-C-based methods on the gut microbiome of humans and animals [21–24,28]. In this this study, we sought to implement meta3C on a human stool sample to link ARGs to their microbial hosts, as well as compare the 3C data generated here to previously published 3C/Hi-C datasets with the aim to optimise analysis methods for 3C/Hi-C data by reducing the impact of spurious intercontig reads.

The proportion of intercontig reads calculated here considerably varied between each dataset, ranging from 0.64% in the K_HiC dataset to 13.74% in P_HiC. In the meta3C libraries that were generated in this study, the fractions of intercontig read pairs were 1.65%. This is lower than expected from the protocol which suggested that 10-15% of the reads will be from cross-linked fragments [31]. However, another study by the same authors using meta3C on human stool samples reported intercontig reads ranging between 1.92% and 14.58% [32]. Additionally, a study that tested the meta3C protocol on a synthetic community also reported that most of their experiments resulted in approximately 1% proximity ligation read rate [33], which suggests that it may be challenging to generate high levels of crosslinks using the original meta3C protocol and that additional enrichment, as in the Hi-C protocol, may be required.

The relatively low average number of intercontig reads in the K_HiC dataset was unexpected. After analysing the shotgun metagenomic reads in the same way as the Hi-C reads by mapping them back to the assembly in each sample of K_HiC, the proportion of spurious intercontig reads was higher than the Hi-C intercontig reads, substantiating the hypothesis that true cross-linking had not been achieved for this Hi-C dataset. Our analyses suggest that the Hi-C procedure may not have worked effectively in most of the K_HiC samples. The authors’ claims on widespread HGT between different phyla in the human gut [22], thus needs further validation as other studies indicate that inter-phylum HGT in the human gut microbiome is a rare event [9,34]. These observations also highlighted that background noise introduced by spurious intercontig reads could interfere with the analysis of the intercontig reads in the 3C/Hi-C datasets.

Spurious intercontig reads can be the result of the formation of spurious ligation products between DNA that originated in different hosts during the experimental process of 3C/Hi-C [35]. They can also occur from sequencing errors [36], and as the results in this study show, they are an inherent issue in short-read sequencing, being present in both the shotgun metagenomic sequence datasets and the WGS short-read data analysed here. The issue of spurious intercontig reads has been relatively underappreciated by the previous studies performing 3C-based techniques on the gut microbiome. On the basis of the analyses in this study, it is clear that they have potential to significantly disrupt the interpretation of data by misassigning hosts to functional genes being investigated. Indeed, when analysing Hi-C reads from wastewater samples, Stalder *et al*. [37] found that several clusters of contigs characterised as Firmicutes, Alphaproteobacteria, and Betaproteobacteria were linked by Hi-C reads to the *E. coli* spike-in strain that had been added to the sample. This *E. coli* spike-in was also linked to several ARGs and plasmids that were not present in the spike-in strain, and the authors concluded that these Hi-C links were spurious and likely due to the high abundance of the spike-in strain [37]. The authors suggested that these ARGs and plasmids were probably present in other strains of *E. coli* that were present in the sample. However, this cannot be confirmed without culturing of the sample. Press *et al*. [24] also observed results that are likely caused by spurious intercontig reads, including a *Eubacterium eligens* megaplasmid being linked by Hi-C reads to another large plasmid originating from a species in the Bacteroidetes phylum.

The majority of the original studies that generated the datasets analysed in this manuscript did little to remove spurious intercontig reads during their analysis. Like the analysis pipeline used in this study, most studies removed reads aligning with a low MAPQ and reads mapping to multiple contigs [21,22,24]. Some also required the presence of restriction sites on the contigs being mapped to [21,22]. However, as these studies used restriction enzymes recognising four-nucleotide motifs, these restriction sites could be quite common in the assembly. Notably, Yaffe and Relman [21] did most to reduce spurious intercontig reads from interfering with the data analysis by developing a pipeline that included probabilistic modelling of experimental noise to determine the likelihood of links made using the 3C data being real. This method allowed the detection and removal of thousands of spurious links. Spurious links can also be removed through normalization of Hi-C data based on zero-inflated negative binomial regression frameworks, although this method has not been applied to 3C/Hi-C experiments on the human gut microbiota [38].

The results in this manuscript show that spurious intercontig reads often account for ∼2% of shotgun metagenomic reads, indicating that a considerable fraction of identified intercontig reads in meta3C/Hi-C datasets, even after removal of low-quality mapping, could be spurious reads that do not originate from cross-linked fragments. Intercontig reads from both 3C data and WGS data were more likely to map near to IS elements. This indicates that many spurious intercontig reads could be caused by repeats in the genome leading to fragmentation of the assembly into smaller contigs. Repeat regions in the genome, like IS elements, that are longer than the read length cause breaks in the assembly as the assembly software will not be able to determine which sequences the repeat is between in the genome. This results in fragmented assemblies in which the repeats are represented as separate individual contigs [39]. This is especially an issue for bacteria, as repeat regions are estimated to make up around 5-10% of the total genome [40]. The typical lengths of IS range between 1,000 and 1,750 bp [41], which is longer than read-length and insert sizes used in 3C/Hi-C. It is thus likely that one of the reads of a pair could map to an IS element contig, or even a contig flanked by repeats. This could cause not only spurious intercontig reads, but also false host-associations of contigs during analysis of 3C/Hi-C data, as the same IS elements can be present in different species [41]. This is particularly relevant for ARGs, which are often flanked by IS elements [42]. Furthermore, our results also showed that intercontig reads were much more likely to map within the first or last 500 nt of a contig compared to non-intercontig reads for both the 3C and WGS reads for the spike-ins. The ends of contigs often contain fragments of repeats [43,44]. By filtering out reads that mapped to IS elements and those that map to the first or last 500 nt of a contig, many of the spurious intercontig reads will be removed. Whilst this may also remove some true intercontig reads that originated from cross-linked fragments of DNA, it is an important step to reduce the impact of spurious intercontig reads on host-ARG associations during further analysis.

Our study also highlights the importance of spike-ins, with completely sequenced genomes, in 3C/Hi-C experiments. Here, a spike-in of two strains of *E. coli* and *E. faecium* were added to the stool sample before meta3C. This was the first study to add spike-ins during proximity ligation of a stool sample, although Marbouty *et al*. [28] added meta3C reads post-sequencing from three bacterial species into the mouse faecal meta3C reads before downstream analysis, and a study implementing Hi-C on wastewater used an *E. coli* spike-in strain in one of the samples [37]. Whilst the G_3C spike-ins were useful in analysis of the meta3C data, by providing positive controls for linkage between ARGs and hosts, the strains used may not have been optimal. Both spike-ins were species of bacteria that are commonly found in the human gut microbiome [45,46]. This meant that any *E. coli* or *E. faecium* strains that were naturally present in the sample used would have been masked by the spike-in strains, complicating the detection of potential ARG-host links to these species. For future 3C/Hi-C experiments, strains of species that are unlikely to be naturally present in the sample type that is being studied should be considered. Ideally these strains should carry resistance genes on both plasmids and chromosomes to corroborate ARG-host linkages on different replicons.

After filtering out spurious intercontig reads, 87 ARGs were linked to their microbial hosts across the 6 datasets, including 27 in the meta3C data first described in this manuscript. These included 6 ARGs known to be carried on plasmids in two spike-in strains that were added to G_3C, showing that meta3C was able to link ARGs carried on plasmids to chromosomal DNA of their microbial hosts in a human stool sample. A potential limitation of our study is that Kraken2 was used to taxonomically classify the linked contigs to determine the hosts of the ARGs. This tool classifies sequences by finding the lowest common ancestor (LCA) of genomes containing an exact match to each *k-mer* in the sequence [47]. Kraken2 relies heavily on correct annotations in the database being used, which is especially a problem when the query contigs differ greatly from sequences in the database [48]. The hosts of some ARGs were very likely misclassified, including the linkage of *tet*(Q) to the fungal genus *Saccharomyces* in the M_3C dataset, *lnu*(P) linking to *Kosmotoga*, a thermophile found in hydrothermal systems in the ocean [49] in Y_3C_A, and *bla*_TEM-116_ in Y_3C_B linking to human mastadenovirus C. It should be noted that these links represented less than 3% of the ARG-host cross-links for those genes. Other 3C/Hi-C studies have used binning methods to improve the reliability of the gene-host link, as this will link genes to a group of contigs rather than just one, which could reduce the chance of misclassifying the host [20]. Classifying these metagenomic assembled genomes (MAGs) often uses phylogenetic trees of multiple marker genes, and whilst this is a well-established method, interpreting the resulting phylogeny and taxonomically classifying the MAGs still has the limitations of needing an accurate reference database [3,48].

Nevertheless, the results of this study showed that ARGs were widespread amongst different microbial hosts, including in many known commensals in the gut microbiome. Genes that were present in multiple datasets showed similar hosts across the datasets. The genes *tet*(Q), *tet*(X), and *erm*(F) were associated with the genera *Alistipes* and *Bacteroides* in nearly all datasets in which these genes occurred. The *tet*(Q), *tet*(X), and *erm*(F) genes are known to be prevalent amongst *Bacteroides* species, and commonly occur together in the same strains, along with the presence of a conjugative transposon [50]. These genes have also been observed in an *Alistipes* strain isolated from the chicken gut [51]. The beta-lactamases *cfxA3* and *cfxA5* were exclusively linked to contigs assigned to the *Bacteroides* genus, where these genes are known to be prevalent [52]. Other genes were widespread and were linked to multiple hosts, including various tetracycline resistance genes, which are highly prevalent and widespread in the human gut microbiota [7,53,54]. Novel observations include the linkage of the vancomycin resistance genes *vanHDX* to the genus *Eubacterium* in the Y_3C_B dataset. This gene has not been observed in *Eubacterium* previously, although *vanD* has been found in several other Eubacteriales, including *Ruminococcus* and *Blautia* [55,56]. Notably, VanD-type glycopeptide resistance genes in gut commensals can be transferred to the opportunistic pathogen *E. faecium*, complicating therapy of infections caused by this species [57].

Overall, the findings in this chapter demonstrate that 3C/Hi-C data contains a substantial background noise from spurious intercontig reads, that could confound host-ARG associations during analysis. Several steps should be taken to reduce the impact of these spurious intercontig reads, including discarding reads that map near to the ends of a contig, removing reads mapping to IS element contigs, and requiring at least five unique intercontig read pairs to link two contigs together. In addition, the use of spike-ins as a control for the efficacy of the cross-linking step in 3C/Hi-C is recommended to ensure the validity of the data.

## Materials and methods

### Stool sample and ethics

The stool sample used to create a meta3C library was obtained from a patient with inflammatory bowel disease, an illness that is associated with higher levels of ARGs in the gut microbiota [58]. Ethical approval for this study has been obtained from the Bradford Leeds Research Ethics Committee (REC 16/YH/0100). The stool sample was divided into ∼500 mg aliquots and stored at -80°C until use.

### Strains

Strains used for spike-in were stored as stocks with 15% glycerol (v/v) at -80°C. *E. coli* E3090 [26] was grown in lysogeny broth (LB) (Sigma-Aldrich), and *E. faecium* E745 [27] was grown in brain heart infusion (BHI) broth (Sigma-Aldrich), both at 37°C with shaking at 200 rpm. To determine viable counts in an overnight broth culture, 10-fold dilutions were made in phosphate-buffered saline (PBS), spread-plated onto the respective agars and incubated at 37°C for 24 h.

### Estimation of the abundance of bacterial cells in stool

To estimate the number of bacterial cells per gram of stool, the copy number of the 16S rRNA gene in the stool sample was estimated as previously described [59]. In short, amplicons (111 nt), generated with primers targeting the V6 region of 16S rRNA gene (5’-CAACGCGARGAACCTTACC-3’ and 5’-ACAACACGAGCTGACGAC-3’ [60]), of *E. coli* MG1655 were cloned into the pJET1.2 cloning vector (Thermo Scientific). The number of 16S rRNA gene copies in stool were then determined using quantitative PCR with the above primers for a concentration range of the pJET1.2-16S construct and the DNA isolated from 400 mg stool, using the FastDNA™ Spin Kit for Soil (MP Biomedicals). We used 2× Luna® Universal qPCR Master Mix (New England Biolabs [NEB]) for quantitative PCR in a volume of 20 µL and primer concentrations of 250 nM each for the forward and reverse primers. The qPCR was then run, in triplicate, on a Bio-Rad CFX Connect™ Real-Time PCR Detection System, following the Luna® protocol. To estimate the number of bacterial cells in the stool sample, the 16S rRNA copy number was divided by 3.82, the average 16S rRNA gene copy number in bacteria [61].

### meta3C

Meta3C was carried out following the protocol from [31], summarised below. Before cross-linking was performed on the stool sample, a spike-in of *E. coli* E3090 and *E. faecium* E745 was added to a final concentration of 1% (0.5% each), calculated using the viable counts of overnight cultures and the estimated number of cells/g of stool, as described above.

Approximately 250 mg of stool was added to 25 mL of PBS with 5% methanol-free formaldehyde (Sigma-Aldrich). After resuspension by vortexing for 30 s, the stool was incubated for 30 min at room temperature (RT) with shaking (250 rpm), followed by 30 min at 4°C under gentle agitation (33 rpm using a roller mixer). Glycine (Fisher Scientific) was then added to a final concentration of 420 mM to quench remaining formaldehyde and incubated for 5 min at RT with moderate shaking (120 rpm), followed by 15 min at 4°C under gentle agitation. The sample was then centrifuged at 4,800 × g for 10 min at 4°C. The pellet was washed with sterile distilled water and resuspended in 4 mL of 1× TE (Tris/EDTA) buffer pH 8.3 (Sigma-Aldrich) supplemented with cOmplete™ mini EDTA-free protease inhibitor (Roche Diagnostics). The suspended pellet was then transferred to four Lysing Matrix E tubes (MP Biomedicals) and run on the FastPrep-24 bead-beater (MP Biomedicals) for three cycles of 8.0 m/s for 20 s, off for 30 s. This run of three cycles was repeated three times, with cooling of the tubes on ice for 5 min between each run. After transfer of the lysate to 15 mL tubes, sodium dodecyl sulphate (SDS) (National Diagnostics) was added to the samples to a final concentration of 0.5% and, after mixing by inversion, the tubes were incubated for 20 min at 65°C, then cooled on ice. The DNA was then digested using 1000 units of either MluCI or HpaII in 1× NEB1 digestion buffer (NEB) and 1% Triton X-100 (Sigma-Aldrich) for 3 h at 37°C. The digestion reaction mixes were centrifuged at 16,000 × g for 20 min at 4°C, and each pellet was resuspended in 500 µL of cold sterile distilled water. Separate ligation reactions (total volume 16 mL) were prepared for the MluCI and HpaII-digested DNA with mixes were prepared containing 1× ligation buffer (50 mM Tris-HCl pH 7.4 (Jena Bioscience), 10 mM MgCl_2_ (Sigma-Aldrich), 10 mM DTT (Roche Diagnostics)) and 0.1 mg/mL bovine serum albumin (BSA) (Sigma-Aldrich). To start the ligation reaction, adenosine triphosphate (ATP; Roche Diagnostics), to a final concentration of 1 mM, and 250 U of T4 DNA ligase (NEB) were added to the ligation reaction tubes, which were then incubated at 16°C for 4 hours. Reversal of the cross-links was then carried out by the addition of 200 µL 0.5 M EDTA, 200 µL 10% SDS, and 100 µL 20 mg/mL proteinase K to the ligation reactions, followed by overnight incubation at 60°C. DNA was then further purified using extraction with phenol-chloroform-isoamyl alcohol and precipitation with isopropanol and ethanol. The purified DNA pellets were resuspended in 60 µL Tris-HCl pH 7.5 with 0.8 mg/mL RNAse A (Qiagen) and incubated at 37°C for 30 min. The quality and quantity of DNA was assessed by performing gel electrophoresis and the Qubit dsDNA BR Assay Kit (Thermo Scientific), respectively. DNA was stored at -20°C until library preparation.

Meta3C sequencing libraries were generated with the NEBNext^®^ Ultra™ II FS DNA Library Prep Kit for Illumina (NEB catalogue number #E6177) following the manufacturer’s protocol with barcoding of the MluCI and HpaII libraries with the NEBNext^®^ Multiplex Oligos for Illuminia^®^ (NEB #E7335). The libraries were quantified on a 2200 TapeStation system (Agilent) using the High Sensitivity D5000 reagents and ScreenTape (Agilent) as per the manufacturer’s protocol to ensure fragmentation ranging between 300-1000 bp. Prepared sequencing libraries were the sequenced by Genomics Birmingham on an Illumina NextSeq 2×150 paired-end platform using a Mid Output Kit v2.5 (300 cycles) (Illumina) with a 1% PhiX spike-in. This dataset is named G_3C in this publication and the short read data are available in the European Nucleotide Archive (ENA), accession number PRJNA879122.

### Analysis of 3C/Hi-C datasets

Reads from published 3C/Hi-C gut microbiome studies were downloaded from the short read archive (SRA) (<u>Table</u>) using the `fastq-dump` of the SRA-Toolkit [62] with the `--split-files` option.

We used identical workflows for G3_C and the downloaded datasets. All Bash and R scripts used in this workflow are available at https://github.com/gregmcc97/3C-HiC_analysis. Duplicate reads were removed using PrinSeq-lite [63]. Reads were then quality filtered (`--nextseq-trim=20` or `-q 20` and min length 60 nt) and had adapter sequences removed using CutAdapt v2.5 [64]. Human sequences were removed with Bowtie2 v2.3.4.1 [65], BEDtools v2.25.0 [66], and Samtools v0.1.19 [67] using the GRCh38.p13 human reference genome (or the GRCm38.p6 mouse reference genome for the M_3C dataset) from the National Centre for Biotechnology Information (NCBI) [68]. The remaining high-quality, non-human, unique, paired reads were then assembled using MEGAHIT v1.1.3 [69] using default parameters and filtering out contigs shorter than 1 kb (`--min-contig-len 1000`). The taxonomic profile of the processed reads was generated using MetaPhlAn3 v3.0 (`--unknown-estimation -–add-viruses`) [29].

ARGs were identified using ABRicate v0.9.8 (https://github.com/tseemann/abricate) with the ResFinder database [70] (≥75% coverage, ≥95% identity). To calculate the abundance of the ARGs, they were first extracted from their contigs and CoverM v0.4.0 (https://github.com/wwood/CoverM) was used to calculate the number of reads mapping to each ARG. The number of mapped reads was then used to calculate the reads per kilobase per million mapped reads (RPKM) using the following formula:

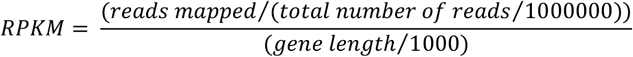

The first 50 bp of the 3C/Hi-C reads were mapped to their respective assemblies using the Burrows-Wheeler Alignment Tool v0.7.12 [71] using the aln and sampe sub-commands. The aligned reads were then filtered to remove those with a mapping quality <20 using Samtools. Read pairs where each mate of the pair mapped to a different contig (intercontig reads) were then identified using Samtools (`view -F 14`) to filter out reads in the SAM file that mapped in a proper pair, were unmapped, or had an unmapped mate, followed by the Unix “grep” command to remove reads in the SAM file that mapped to the same contig as their mate (`-grep -v “=“`).

### Analysis of G_3C reads mapping to spike-ins

The complete genome sequences of the *E. coli* E3090 (assembled as described in [26]) and *E. faecium* E745 (downloaded from NCBI, accession GCA_001750885.1) spike-ins were annotated using Prokka [72]. WGS reads (ENA accession numbers: *E. coli* E3090: ERX2620237; *E. faecium* E745: SRS15053183) and 3C reads were then mapped to the genomes and an R script (available at https://github.com/gregmcc97/3C-HiC_analysis) was used to identify the annotated region of the genome being mapped to by each read. From the output file, products labelled as “NA” were assigned as intergenic regions. Products labelled as “*IS*” or “*transposase*” were assigned as IS elements. Products labelled as

“*hypothetical_protein*” or “*product=putative protein*” were assigned as genes with unknown functions. Remaining products labelled as “*gene*”, “*locus_tag*”, “*db_xref*”, “*protein*”, “*note*”, or “*product*” were assigned, using a bash script, as genes with predicted functions. To calculate the proportion of reads that map within the first or last 500 nt of a contig, a bash script was used that mapped coordinates in the SAM mapping file and the contig lengths in the assembly.

### Filtering of spurious intercontig reads

A bash script (available at https://github.com/gregmcc97/3C-HiC_analysis) was written to remove intercontig reads that mapped within the first or last 500 nt of a contig. Further filtering was carried out after intercontig reads linking contigs to ARG contigs were identified (see below).

### Linking ARGs to their microbial hosts

3C/Hi-C intercontig reads where one mate mapped to a contig carrying an ARG gene were identified to generate a list of linked contigs for each ARG contig. These lists were then filtered so that only contigs that linked at least five times to an ARG contig were kept. Additionally, and to remove potential false cross-links from contigs that contain IS elements, IS elements in the assembly were identified using ABRicate with the ISfinder [73] database (≥60% coverage, ≥99% identity) and these were removed from the lists of contigs linked to ARGs.

Remaining contigs for each ARG were then taxonomically classified using Kraken2 v2.0.8 [74] using the prebuilt kraken2-microbial database (https://lomanlab.github.io/mockcommunity/mc_databases.html). The contigs were also mapped to NCBI’s nucleotide (nt) database using BLASTN v2.2.31 [75]. Links to contigs that aligned with 99% identity to known plasmid sequences using BLAST were removed. Pheatmap (https://github.com/raivokolde/pheatmap) was used to create a heatmap of the ARG-host associations.

## Acknowledgements

The authors thank Dr Steven Dunn for helpful discussions on bioinformatics. GEM was supported by the MRC IMPACT DTP at the University of Birmingham (MR/N013913/1). WvS was supported by a Royal Society Wolfson Research Merit Award (WM160092). Sample collection was supported by a University Hospitals Birmingham Charity Award to THI.

## Author contributions

SAK and WvS designed the study. GEM performed the experimental work and analyses. AER collected stool samples. THI and MNQ were responsible for obtaining ethical approval for this study. GEM wrote the first draft of the manuscript, with further editing by WvS, AER and SAK. All authors reviewed and approved the final manuscript.

