## Supplementary Table 1 for "Noise reduction strategies in metagenomic chromosome confirmation capture to link antibiotic resistance genes to microbial hosts"

### Supplementary Table 1. Read counts during meta3C/Hi-C analysis

| Dataset | G_3C | M_3C | Y_3C | | D_HiC | P_HiC | K_HiC* |
|---|---|---|---|---|---|---|---|
| | | | Y_3C_A | Y_3C_B | | | |
| **Processed reads** | 198,493,086 | 366,961,002 | 2,921,579,828 | 1,239,950,680 | 133,509,800 | 157,755,162 | 37,984,239 |
| **Reads mapped (MAPQ>20)** | 181,467,148 | 278,726,053 | 2,868,601,794 | 1,155,939,113 | 108,556,752 | 124,877,330 | 32,412,518 |
| **Percentage mapped** | 91.42% | 75.96% | 98.19% | 93.22% | 81.31% | 79.16% | 85.66% |
| **Intercontig reads** | 3,271,007 | 35,717,451 | 188,322,547 | 94,104,831 | 9,488,683 | 21,679,019 | 192,510 |
| **Percentage intercontig** | 1.65% | 9.73% | 6.45% | 7.59% | 7.11% | 13.74% | 0.64% |
| **Filtered intercontig reads** | 1,574,468 | 28,773,234 | 90,197,910 | 63,855,164 | 6,880,255 | 18,917,493 | 53,157 |
| **Percentage intercontig after filtering** | 0.79% | 7.84% | 3.09% | 5.15% | 5.15% | 11.99% | 0.18% |

*for K_HiC, an average of 43 samples is presented in this table; For datasets that used multiple restriction enzymes, numbers presented are a combined total; MAPQ = mapping quality
