## Supplementary Table 2 for "Noise reduction strategies in metagenomic chromosome confirmation capture to link antibiotic resistance genes to microbial hosts"

**Supplementary Table 2. Comparison of spike-in WGS and G_3C reads that map to spike-in genomes**

| **Spike-in** | ***E. coli* E3090** | | ***E. faecium* E745** | |
|---|---|---|---|---|
| **Dataset** | **G_3C** | **WGS** | **G_3C** | **WGS** |
| **Total reads** | 14,497,782 | 1,284,538 | 8,170,430 | 3,333,334 |
| **Reads mapped to G_3C assembly (MAPQ>20)** | 14,237,338 | 1,256,272 | 7,764,542 | 3,175,581 |
| **%mapped to G_3C assembly** | 98.20% | 97.80% | 95.03% | 95.27% |
| **Intercontig reads** | 141,473 | 10,080 | 81,191 | 78,106 |
| **%intercontig reads** | 0.98% | 0.78% | 0.99% | 2.34% |

WGS = whole genome sequencing; MAPQ = mapping quality; nt = nucleotides
