## Supplementary Figure 1 for "Noise reduction strategies in metagenomic chromosome confirmation capture to link antibiotic resistance genes to microbial hosts"

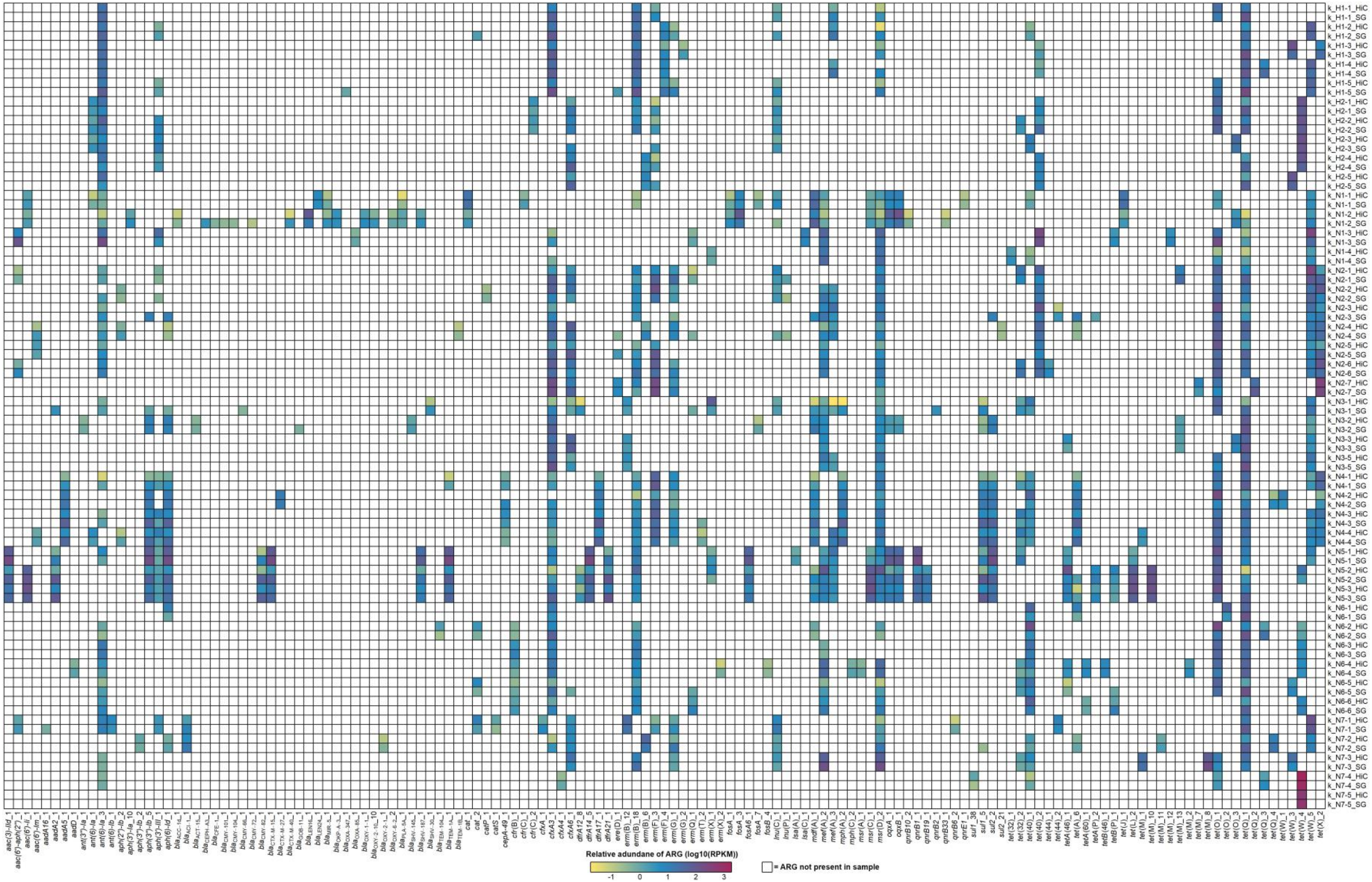


**Supplementary Figure 1. ARG abundance in the K_HiC dataset.**

ARG sequences from the assemblies were isolated, and the reads from each dataset were mapped to the ARGs (columns). The relative abundance was calculated as reads per kilobase per million mapped reads (RPKM). White cells mean the ARG was not present, and coloured cells show that the ARG was present, with the colour relating to the relative abundance of the ARG within that set of reads (log(10) transformed RPKM values). Different datasets are separated by gaps in the heatmap. The individual datasets show RPKM of the shotgun reads (*_SG) or Hi-C reads (*_HiC) mapping to ARGs identified in the shotgun metagenomic assembly.
