## Supplementary Figure 2 for "Noise reduction strategies in metagenomic chromosome confirmation capture to link antibiotic resistance genes to microbial hosts"

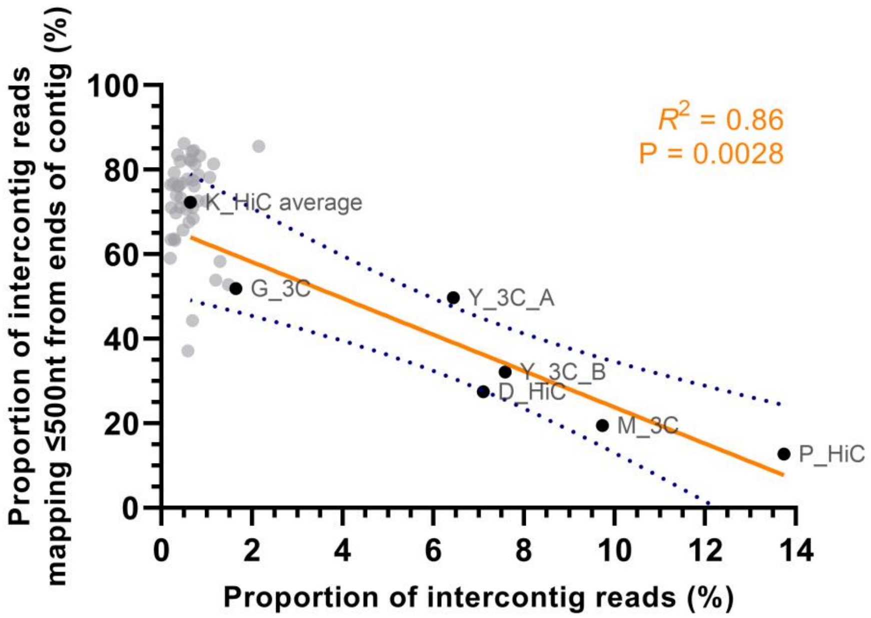


**Supplementary Figure 2. The proportion of identified intercontig reads vs the proportion of intercontig reads mapping within the first or last 500 nucleotides (nt) of a contig for all 3C/Hi C datasets.**

Each point represents a different 3C/Hi-C sample (labelled). Unlabelled grey points are the individual samples in the K_HiC dataset. Slope calculated via linear regression analysis showing a statistically significant correlation (P = 0.0028, Spearman correlation). Blue dotted lines indicate the 95% confidence interval.
